# Limitations of grouping subjects based on biological sex (males versus females) and a new approach: insights from intra-nucleus accumbens core dopamine-induced psychostimulant activity

**DOI:** 10.1101/2024.10.22.619725

**Authors:** Martin O. Job

## Abstract

**Background:** In the field of substance use disorder research, sex-as-a-biological-variable (SABV) is employed to determine the mechanisms governing sex differences. Based on our recently developed MISSING (Mapping Intrinsic Sex Similarities as an Integral quality of Normalized Groups) model, we hypothesized that grouping subjects by biological sex does not represent the most effective way to group behavioral data objectively. To test our hypothesis, we conducted experiments to compare the psychostimulant effect of intra-nucleus accumbens (NAc) dopamine on groups based on 1) biological sex (current model) and 2) behavioral clustering (MISSING model) for effectiveness in identifying groups of subjects that a) are distinct with regards to behavioral variables, and b) confirm NAc dopamine neurochemical expression/activity topography (NEAT).

**Methods:** For the current model, we separated subjects (n = 37 Sprague Dawley rats, male n = 20, female n = 17) by biological sex prior to all assessments. For the MISSING model, we conducted normal mixtures clustering of baseline activity, dopamine activity (as distance traveled in cm over 60 min) and dopamine activity normalized-to-baseline activity (NBA) of all subjects to identify behavioral clusters.

**Results:** Separating groups by biological sex revealed groups (males and females) that were not clearly distinct with regards to behavioral variables and do not confirm NAc dopamine NEAT. Separating groups using the MISSING model revealed groups (behavioral clusters) that were clearly distinct with regards to behavior and confirm NAc dopamine NEAT.

**Conclusions:** Our results reveal the limitations of grouping subjects based on biological sex. We discuss a new approach.

## Introduction

Psychostimulants increase locomotor activity in males and females in preclinical models. It is generally assumed that females are more responsive to these drugs than males (Cailhol and Mormède, 1999; Chin et al., 2002; Craft and Stratmann, 1996; Festa et al., 2004; Gaines et al., 2022; Gogos et al., 2017; Harrod et al., 2005; Job et al., 2014a; King et al., 2021; Kuhn et al., 2001; Leong et al., 2016; McDougall et al., 2020, 2015; Milesi-Hallé et al., 2007, 2005; Ohia-Nwoko et al., 2017; Parylak et al., 2008; Robison et al., 2017; Schindler et al., 2002; Schindler and Carmona, 2002; Segarra et al., 2010; Sershen et al., 1998; Siegal and Dow-Edwards, 2009; Šlamberová et al., 2013, 2011; Thomsen and Caine, 2011; Van Haaren and Meyer, 1991; Van Swearingen et al., 2013; Wissman et al., 2011; Zhou et al., 2012), *but this is not always the case* (Cailhol and Mormède, 1999; Carroll et al., 2007; Collins et al., 2015; Gaines et al., 2022; Martz et al., 2023; McDougall et al., 2018; Ramos et al., 2020; Risca et al., 2020; Thomsen and Caine, 2011; Zombeck et al., 2010). There are also sex differences in baseline locomotor activity, with females showing greater activity than males (Cailhol and Mormède, 1999; Chin et al., 2002; Milesi-Hallé et al., 2007, 2005; Ohia-Nwoko et al., 2017; Ramos et al., 2020; Segarra et al., 2010; Sershen et al., 1998; Simpson et al., 2012; Simpson and Kelly, 2012; Van Haaren and Meyer, 1991; Van Swearingen et al., 2013; Zombeck et al., 2010), *but not always* (Ohia-Nwoko et al., 2017; Robison et al., 2017; Schindler et al., 2002; Schindler and Carmona, 2002). From the above, we can infer that there are contexts where we can detect *both* sex differences and sex similarities in baseline and psychostimulant-induced locomotor activity. In the field, these similarities/differences generally thought to be linked to ‘sex as a biological variable’ (SABV).

But biological sex *per se* can be a very complex variable especially because males and females do not represent behaviorally-homogenous populations. Indeed, when we account for this heterogeneity, we can observe sex differences, or not, depending on what within-sex population groups we compare (Brown et al., 2015; Carreira et al., 2017; Carroll et al., 2007; Davis et al., 2008). This suggests that without accounting for within-sex heterogeneity, we may not be getting the complete picture as it relates to sex differences/similarities.

In the current model in the field, subjects are first identified/separated by biological sex *before* behavioral assessments are observed/compared (Figure 1A). The Mapping Intrinsic Sex Similarities as an Integral quality of Normalized Groups (MISSING) model (see Figure 1B, compared to the current model in Figure 1A) employs objective clustering analysis of all subjects, *irrespective of biological sex*, to identify individuals that are similar/different – if males and females represent behaviorally distinct groups they should separate into different clusters.

**Figure 1:**
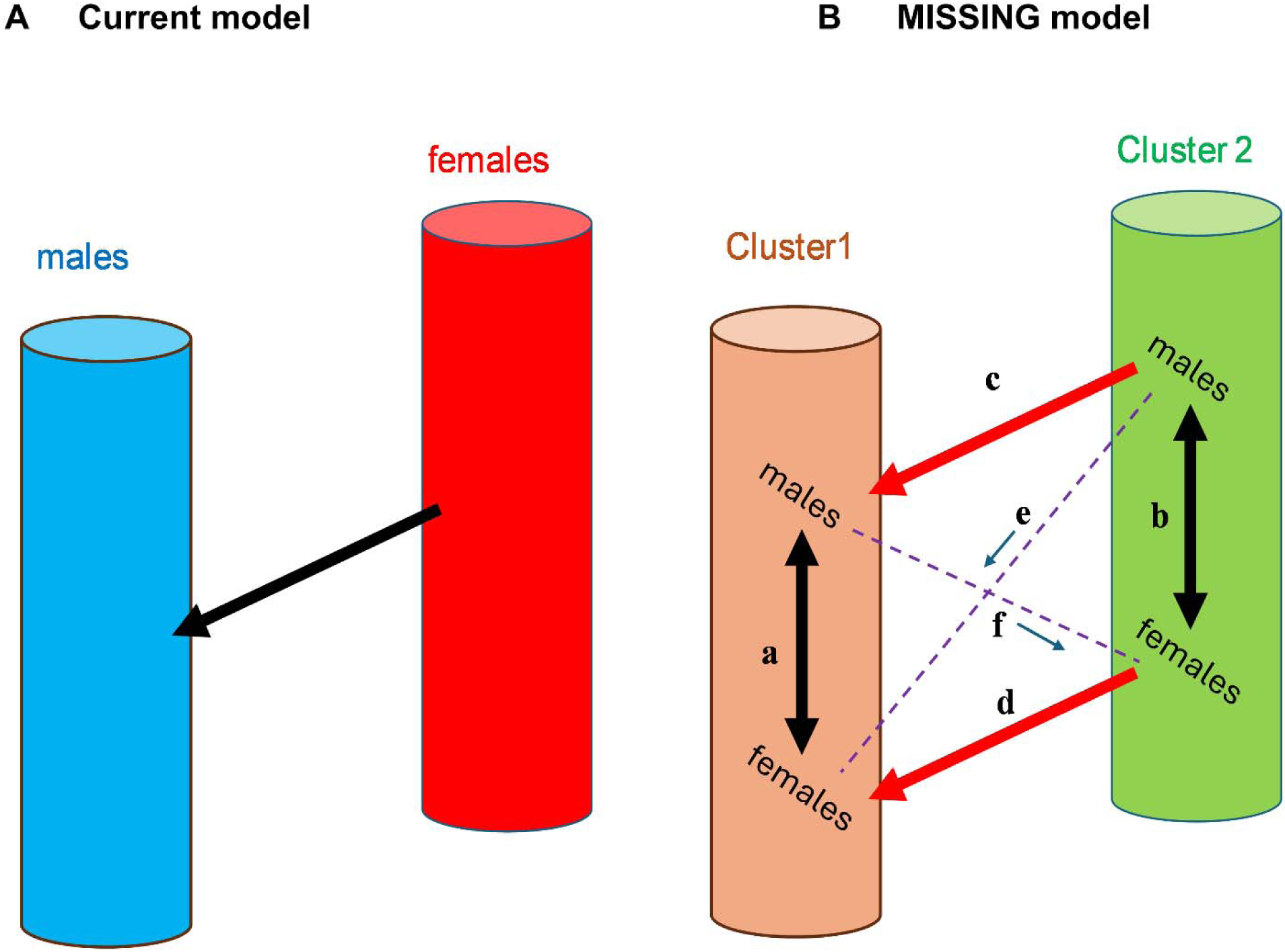
Defining the current and the MISSING models. Fig A represents the current model in which subjects are grouped based on biological sex per se prior to behavioral assessments. Essentially the experimenter knows which subjects are males and females and assumes that they are already different and then tries to understand how they are different, or not, using behavioral tests. Fig B represents the Mapping Intrinsic Sex Similarities as an Integral quality of Normalized Groups (MISSING) model. Here, the experimenter is deliberately blinded to the biological sex of the subject and tries to objectively identify what behavioral cluster the individual belongs to, with observation of sex differences/similarities assessed only after this behavioral cluster has been identified. In Fig B, within a cluster, males and females are similar (a, b) and males and females in a cluster are significantly different with regards to behavioral typology than males and females in another cluster (c, d). MISSING model proposes that sex differences, if any, are mostly related to cluster differences (e and/or f are related to c and/or d), not biological sex per se. Fig A groups by biological sex while Fig B groups via objective clustering. Our hypothesis is that grouping subjects by biological sex (Fig A) does not represent the most effective way to group behavioral data **objectively.**

This novel MISSING model was recently validated for cocaine-induced locomotor activity (Tigano and Job, 2024) and it revealed that the most distinct groups were not between sexes but between behavioral clusters.

Cocaine, like other psychostimulants, exerts its locomotor effects partly via increasing dopamine activity in a limbic region of the brain termed the nucleus accumbens (NAc) (Kelly and Iversen, 1976; Sharp et al., 1987). Direct injection of dopamine into the NAc mimics the effects on locomotor activity of intra-NAc injections of psychostimulants (Campbell et al., 1997; Delfs et al., 1990; Essman et al., 1993; Heidbreder and Feldon, 1998; Ikemoto, 2002; Ikemoto and Witkin, 2003; Job et al., 2014b; Swanson et al., 1997). The neurochemical mechanisms governing sex differences in psychostimulant effects are not fully understood, but are thought to have something to do with biological sex-specific differences in the effects of these drugs on NAc dopamine activity (Becker, 1999). The NAc is a heterogeneous structure that is divided into shell and core subregions (Meredith et al., 2008; Pennartz et al., 1994; Salgado and Kaplitt, 2015; van Dongen et al., 2005; Zahm, 1999). Of these NAc regions (shell and core), the NAc core is the more sexually dimorphic (Table 1). Thus, to effectively assess sex differences in intra-NAc dopamine activity it is important to focus on the NAc core (Table 1). As the NAc is topographically-organized with regards to the dopamine system (Table 2), so also is the NAc core, see (Jiang et al., 2014; Lee et al., 2014; Pennartz et al., 1994; Prast et al., 2014b, 2014a; van Dongen et al., 2008, 2005). Thus, dopamine injection into the NAc should exhibit a neurochemical expression/activity topography (NEAT). Based on the expression gradient of endogenous dopamine expression (row 1, Table 2), a NEAT analysis for the effects of exogenous dopamine injected into the NAc should reveal an activity gradient along the anterior-posterior (AP), medial-lateral (ML) and dorsal-ventral (DV) axis.

**Table 1:** The nucleus accumbens core is more sexually dimorphic than the nucleus accumbens shell.

| Observation | NAc core | Sex differences<br>NAc shell | References |
| --- | --- | --- | --- |
| MSNs: Dendritic spine density/ spine heads | yes | no | (Forlano and Woolley, 2010; Wissman et al., 2012, 2011) |
| Ketamine-induced increases in MSNs dendritic spine density | yes | no | (Strong et al., 2017) |
| Sensitivity of MSNs to estrogen-mediated regulation of excitatory inputs | yes | no | (Cao et al., 2018, 2016; Peterson et al., 2016, 2015; Proaño et al., 2020, 2018; Proaño and Meitzen, 2020; Willett et al., 2016) |
| Ovariectomy-mediated decreases in [<sup>3</sup>H] quinpirole specific binding (D2R binding) | yes | no | (Le Saux et al., 2006) |
| Oxytocin-mediated increase in extracellular dopamine release following cocaine experience | yes | no | (Logan et al., 2022) |
| Withdrawal from binge-pattern cocaine administration: δ-opioid receptor trafficking | yes | no | (Ambrose-Lanci et al., 2008) |
| Dopamine release/reuptake | yes | no | (Wallace et al., 2022) |
| Effects of opioids | yes | no | (Martin et al., 2015; Mousavi et al., 2007; Wang et al., 2014) |
| D1-D2 heteromers | yes | no | (Hasbi et al., 2020a) |
| Nicotine-mediated expression of Substance P | yes | no | (Pittenger et al., 2016) |

**Table 2:** Expression topography for dopamine system components in the nucleus accumbens.

| DA system component | Anterior-to-Posterior (AP) | Medial-to-Lateral (ML) | Dorsal-to-Ventral (DV) |
| --- | --- | --- | --- |
| DA content | Decreases (Beal and Martin, 1985; Lee et al., 2014; Widmann and Sperk, 1986) | Decreases (Beal and Martin, 1985; Deutch and Cameron, 1992; Lee et al., 2014) | Decreases (Beal and Martin, 1985; Deutch and Cameron, 1992; Widmann and Sperk, 1986) |
| D1 receptor | Decreases (Bardo and Hammer, 1991; Boyson et al., 1986; Richfield et al., 1987) | Unchanged (Berendse and Richfield, 1993)/increases (Savasta et al., 1986)/decreases (Bardo and Hammer, 1991) | Unchanged (Savasta et al., 1986) / increases (Berendse and Richfield, 1993) |
| D2 receptor | Decreases (Alakurtti et al., 2013; Bardo and Hammer, 1991; Boyson et al., 1986; Brock et al., 1992; Jongen-Rêlo et al., 1995; Richfield et al., 1987) | Increases (Allin et al., 1989; Bardo and Hammer, 1991) | Decreases (Allin et al., 1989) |
| DA Transporter | Decreases (Graybiel and Moratalla, 1989; Tassin et al., 1976) | Increases (Freed et al., 1995; Graybiel and Moratalla, 1989; Siciliano et al., 2014) | Decreases (Calipari et al., 2012; Graybiel and Moratalla, 1989; Marshall et al., 1990; Siciliano et al., 2014) |
| D1-D2 receptor heteromer | Decreases (Hasbi et al., 2020b) | Unchanged (Hasbi et al., 2020b)/ increases (Hasbi et al., 2020a) | Increases (Hasbi et al., 2020b) |

### Our hypothesis was that grouping subjects by biological sex does not represent the most objective way to group behavioral data

To test our hypothesis, we conducted experiments to compare grouping via biological sex with grouping via behavioral clustering to determine which was more effective in identifying groups of subjects that are distinct with regards to 1) behavioral variables, and 2) NAc dopamine neurochemical expression/activity topography (NEAT). Our methods, results, discussion and conclusions are below.

## Methods and Materials

### Animals

A total of 37 (males n = 20, females n = 17) adult Sprague Dawley rats (20 males and 17 females), weighing between 225–275 g at the time of purchase (Charles River Inc., Wilmington, MA), were used for the behavioral studies. The rats had access to rat chow and water *ad libitum* and maintained on a 12-hour light: dark cycle (lights on at 7am). Experiments and animal care were in accordance with the Institute of Animal Care and Use Committee of Emory University and the National Institutes of Health Guide for the Care and Use of Laboratory Animals. Two rats of the same sex were housed per cage prior to surgery. The rats were individually housed following surgery to prevent damage to the surgical area by other rat cage occupants.

### Surgery

Surgeries were done as previously described (Job, 2016; Job et al., 2014b, 2013). Briefly, under isoflurane inhalation anesthesia, or with a cocktail of Ketaset (Ketamine HCl, Fort Dodge Animal Health, Iowa, USA) and Dexdomitor (dexmedetomidine HCl, Orion Corporation, Espoo, Finland) injection anesthesia, stereotaxic surgical procedures were employed to implant rats, using stereotaxic surgical techniques, with a bilateral stainless-steel guide cannulae assembly (22 gauge; P1 Technologies; Roanoke, VA) placed 2.0 mm above the NAc core. The target coordinates (from Bregma) for accessing the NAc core were as follows: AP = 1.6 mm, ML = ± 1.5 mm, and DV = −5.7 mm (Paxinos and Watson, 1998). The guide cannulae assembly was secured to the skull using 2–4 stainless steel screws, glue and dental cement. To prevent cannulae blockage, a bilateral obturator extending 0.5mm past the tip of the cannulae was inserted into the guide cannulae. All rats were allowed to recover for at least one week before locomotor activity assessments.

### Habituation

Following recovery from surgery, the animals were prepared for infusions and locomotor activity measurement. One to two days before testing, animals were placed in the photocell cages so that they can explore the testing chambers to habituate them to experimental conditions. This habituation procedure is done for approximately 30 minutes after rats are briefly handled.

### Locomotor Activity Assessment

For details, see (Job, 2016; Job et al., 2014b, 2013), and also see (Tigano and Job, 2024). On the experiment day, rats were placed into the locomotor chambers for 30 min to again habituate to their surroundings before the experiment began. For the experiment, basal locomotor activity for 30 min. After this, rats were removed from the chambers and dopamine (15 µg/0.5 µL bilaterally) was injected into the NAc core and animals were placed back into the chambers for additional recording of locomotor activity for 60 min.

### Perfusions and Histology

Perfusions were done as previously described (Job, 2016; Job et al., 2014b, 2013). After Nissl staining, a stereotaxic brain atlas (Paxinos and Watson, 1998) was used to approximate the bilateral placement of the injector tips for every individual.

### Variables

The variables employed include baseline activity, dopamine activity, dopamine activity NBA – similar to our recent report with cocaine (Tigano and Job, 2024).

Baseline activity was estimated as distance traveled (cm) in 30 min just prior to dopamine injection. Dopamine activity was assessed as distance traveled in 60 min after dopamine injection. dopamine activity normalized to baseline activity (dopamine activity NBA) was calculated as shown in the equation below (see Figure 2)

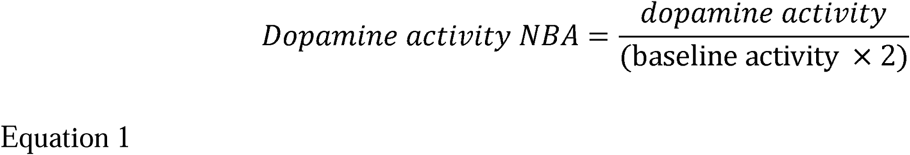

**Figure 2:**
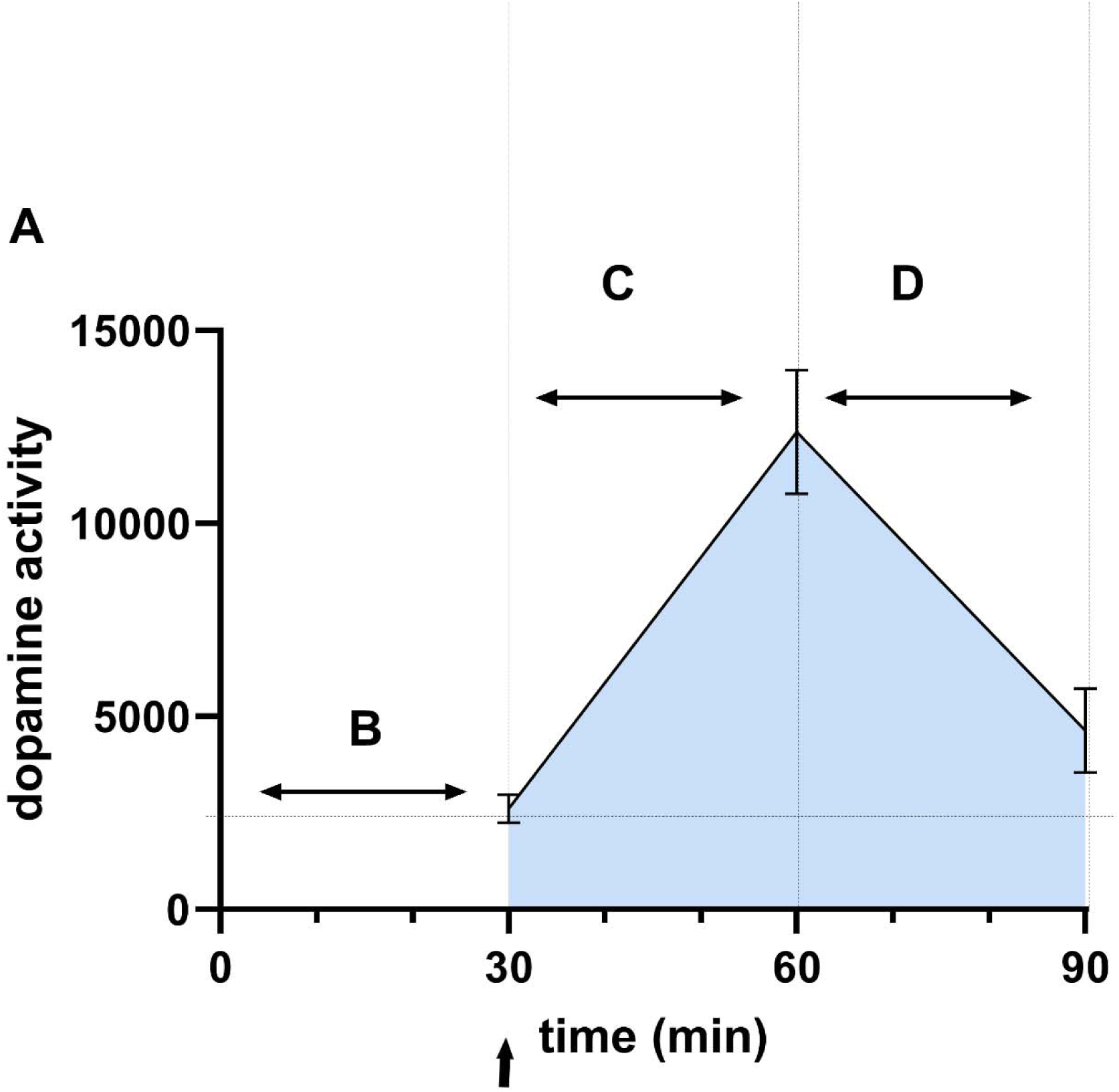
Calculating dopamine activity normalized-to-baseline activity (dopamine activity NBA). The rationale for this variable is that 1. dopamine activity is a change in activity from baseline activity, and 2. while occurring, dopamine activity is always relative to baseline activity (not zero activity). The figure shows how we derived the dopamine activity NBA variable. The x-axis represents time (min), and the y-axis represents the distance traveled in cm (cocaine activity). B represents baseline activity. The arrow at time = 30 min represents the dopamine injection point. The shaded region represents the sum over time (60 min) of the distance traveled and corresponds to the area under the curve. To obtain dopamine activity NBA, we divided total dopamine activity by baseline activity with baseline activity adjusted for time (C-D). Thus, dopamine activity NBA = dopamine activity (60 min) / baseline activity estimated for 60 min (baseline activity × 2), as such dopamine activity NBA = C + D/ B × 2. This is calculated similarly to a recent report (Tigano and Job, 2024).

### Statistical analysis

Statistical analysis: GraphPad Prism v 10 (GraphPad Software, San Diego, CA), SigmaPlot 14.5 (Systat Software Inc., San Jose, CA) and JMP Pro v 17 (SAS Institute Inc., Cary, NC) were employed for statistical analysis. Grubb’s test was used to determine if there were any significant outliers. Data were expressed as mean ± SEM. For the current model, we separated subjects by biological sex and conducted unpaired t-tests to determine if there were sex differences between locomotor activity variables and injection site coordinates. We analyzed the distribution to determine if there were more than one group of subjects in our samples. For the MISSING model, we employed normal clustering of several variables (baseline activity, dopamine activity and dopamine activity NBA) for every individual, regardless of sex, to identify distinct clusters/groups. We confirmed that these clusters/groups were indeed distinct using One-way ANOVA (for variable comparisons) and linear regression analysis (for comparisons of relationships between variables). Thereafter, we employed Two-way ANOVA with factors being SEX (males, females) and cluster to determine if there were main effects of SEX, cluster and SEX × cluster interaction. Statistical significance was set at P < 0.05 for all analyses with Tukey’s post hoc test employed when significance was detected. Using linear regression analysis, we conducted dopamine neurochemical expression/activity topography (NEAT) analysis which involved the determination/comparison of the relationship between injection site coordinates (AP, ML and DV) and dopamine activity for groups separated using 1) biological sex and 2) behavioral clustering.

## Results

### Separation via biological sex: No sex differences in dopamine injection site coordinates

We compared the injection sites between males and females. Every subject had bilateral injection coordinates and as such for n = 20 males, we had 40 injection coordinates and for n = 17 females, we had 34 injection coordinates. For males (n = 40), the mean ± SEM for AP, ML and DV were 1.29 ± 0.07 mm, 1.53 ± 0.04 mm and 7.17 ± 0.07 mm, respectively. For males (n = 34), the mean ± SEM for AP, ML and DV were 1.29 ± 0.09 mm, 1.46 ± 0.03 mm and 7.36 ± 0.08 mm, respectively. The injection site coordinates for all subjects are shown in Figure 3A. Comparisons of males and females using unpaired t-tests revealed no significant differences for AP (P = 0.9356, Figure 3B), ML (P = 0.3922, Figure 3C) and DV (P = 0.3014, Figure 3D) injection site coordinates.

**Figure 3:**
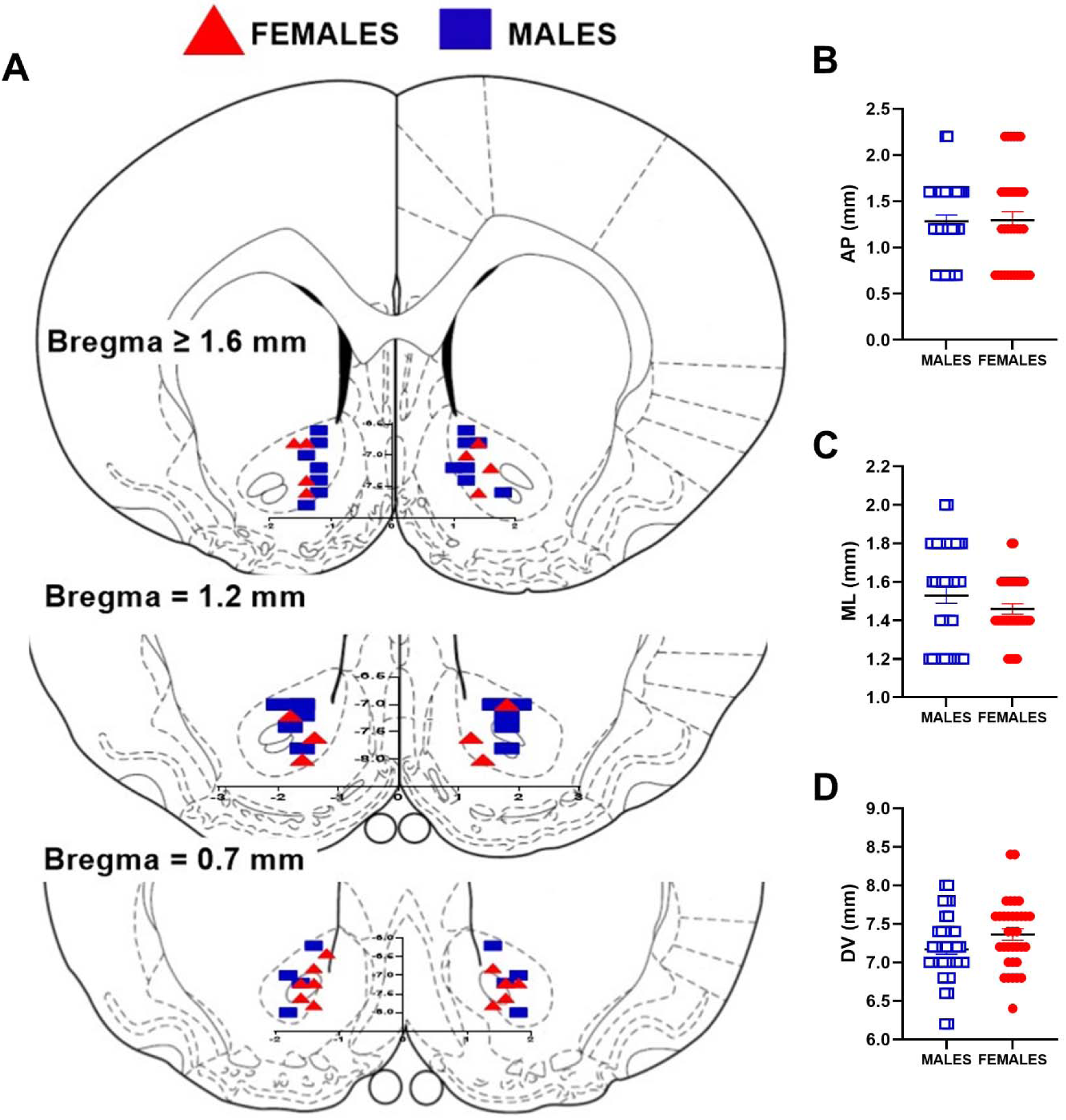
Histology showing placement of injection points bilaterally in male and female Sprague Dawley rats. The brain drawings are from (Paxinos and Watson, 1998) with injection coordinates from bregma (mm). The red triangles represent females while the blue squares represent males (Fig A). We employed unpaired t-tests for a comparison of the injection placements in the anterior-posterior (AP, Fig B), medial-lateral (ML, Fig C) and dorsal-ventral (DV, Fig D) axis of the nucleus accumbens core. For these males and females are shown as blue open squares and red closed circles, respectively. We determined that there were no sex differences with regards to injection tip placements (bilaterally).

### Separation via biological sex: sex differences in dopamine activity

We separated subjects based on biological sex into males and females.

Baseline activity: The mean ± SEM for baseline activity for males (n = 20) and females (n = 17) were 1830 ± 233 and 2601 ± 373 cm, respectively. With regards to baseline activity, Grubb’s test revealed a significant outlier for males, but not for females, and not for all (males and females combined). Unpaired t-tests revealed no significant difference between sexes with regards to baseline activity (P = 0.0793, Figure 4A). Analysis of the data distribution for all males and females suggests the existence of two (2) populations (Figure 4B).

**Figure 4:**
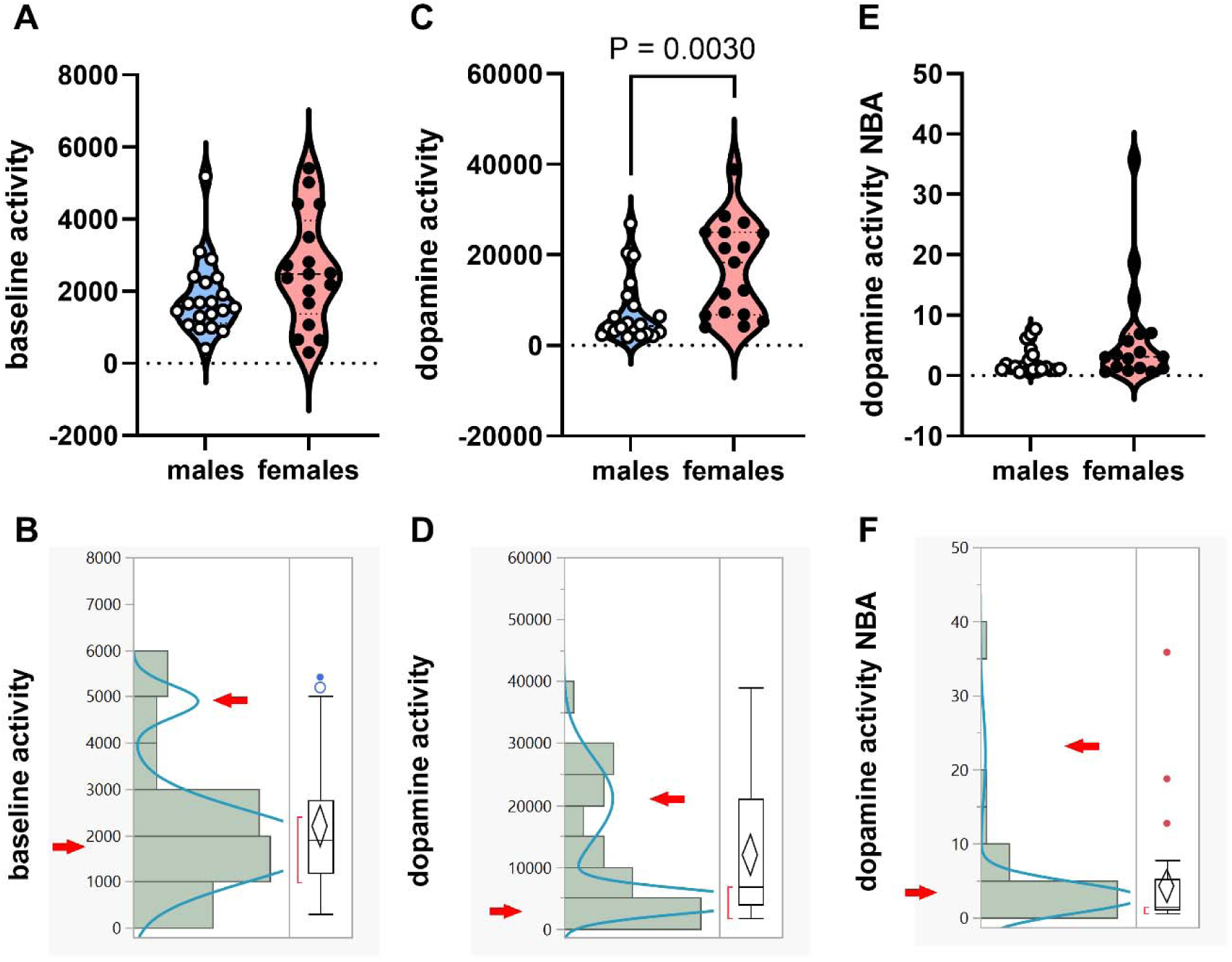
Biological sex differences in dopamine activity: sex differences in dopamine activity complicated by more than one population of subjects in sample: We employed unpaired t-tests to determine if there were sex differences in 1. baseline activity (Fig A), 2. dopamine activity (Fig C) and 3. dopamine activity NBA (Fig E). We analyzed the distribution of data for all subjects (males + females) for baseline activity (Fig B), dopamine activity (Fig D) and dopamine activity NBA (Fig F). There were no sex differences in baseline activity (P > 0.05, Fig A) but the distribution analysis revealed 2 populations (Fig B). There were sex differences in dopamine activity (P < 0.05, Fig C) but the distribution analysis revealed 2 populations (Fig D). There were no sex differences in dopamine activity NBA (P > 0.05, Fig E) but the distribution analysis revealed 2 populations (Fig F). In summary, even when there are sex differences, for example in dopamine activity (Fig C), this finding may be complicated by the presence in the sample set of more than one population of subjects that were not considered in the comparison (Fig D). The arrows shows the outline of the populations within the distribution analysis.

Dopamine activity: The mean ± SEM for dopamine activity for males (n = 20) and females (n = 17) were 7632 ± 1611 and 16995 ± 2556 cm, respectively. For dopamine activity, Grubb’s test revealed no significant outliers for males, females or males plus females. Unpaired t-tests revealed a significant sex difference for dopamine activity (P = 0.0030, Figure 4C). Analysis of the data distribution for all males and females suggests the existence of two (2) populations (Figure 4D).

Dopamine activity NBA: The mean ± SEM for dopamine activity NBA for males (n = 20) and females (n = 17) were 2.34 ± 0.49 and 6.42 ± 2.17, respectively. With regards to dopamine activity, Grubb’s test revealed no significant outlier for males, but did reveal significant outlier(s) for females and for all subjects (males and females combined). Unpaired t-tests revealed no significant difference between sexes with regards to dopamine activity NBA (P = 0.0562, Figure 4E). Analysis of the data distribution for all males and females suggests the existence of two (2) populations (Figure 4F).

### Separation via behavioral clustering (MISSING model)

Because the analysis of distribution suggests there are more than one population in our sample, we considered our sample a mixture and conducted normal mixtures clustering (of all subjects, see MISSING model in Figure 1B) to identify these groups. These groups were separated based on behavioral clustering and not based on biological sex.

Identification of behavioral clusters: Normal mixtures clustering of baseline activity, dopamine activity and dopamine activity NBA for all subjects, irrespective of biological sex (n = 37) revealed 3 clusters which we termed cluster1, cluster2 and cluster3 (Figure 5A-B). Cluster1 and cluster 2 consisted of both males and females while cluster 3 included only females (Figure 5A-B). The group compositions were as follows: cluster1 (n = 23: males n = 16, females n = 7), cluster2 (n = 11: males n = 4, females n = 7) and cluster3 (n = 3: males n = 0, females n = 3).

**Figure 5:**
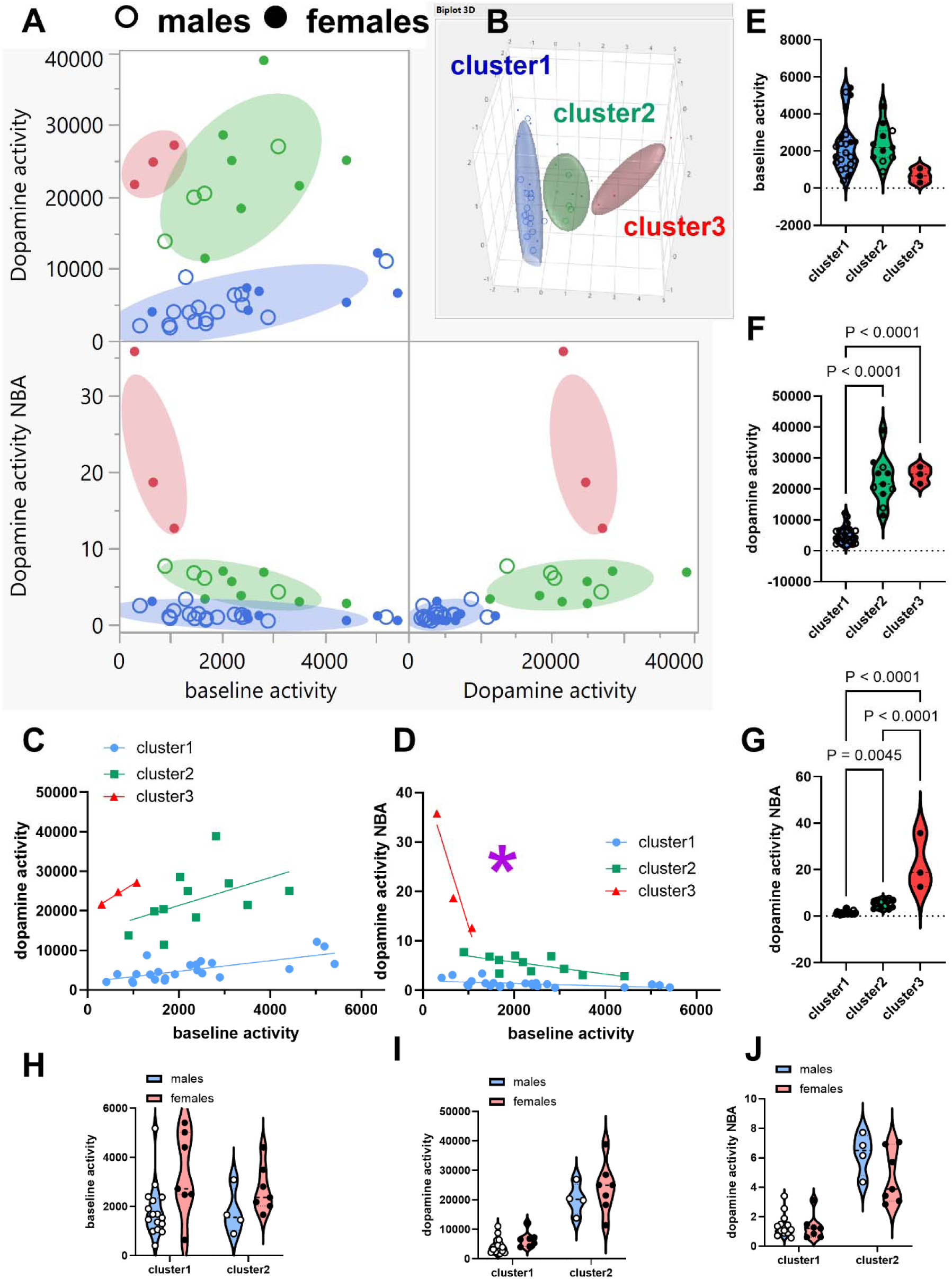
Normal mixtures clustering analysis of baseline activity, dopamine activity and dopamine activity NBA yielded three distinct groups of subjects (males and females). Because of the detection of more than one population of subjects for all variables (Figure 4 B, D and F), we conducted normal mixtures clustering of baseline activity, dopamine activity and dopamine activity NBA for all subjects (n = 37). This analysis yielded three clusters (Fig A-B): cluster1 (n = 23: males n = 16, females n = 7), cluster2 (n = 11: males n = 4, females n = 7) and cluster3 (n = 3: males n = 0, females n = 3). Linear regression revealed no differences between clusters with respect to the slopes of the relationship between baseline activity and dopamine activity (P > 0.05, Fig C). There were significant differences (P < 0.05, Fig D) between the slopes of the relationship between baseline activity and dopamine activity NBA with significant differences when we compared slopes for clusters1 versus cluster3 (P < 0.0001) and cluster2 versus cluster3 (P < 0.0001). One-way ANOVA revealed no significant differences between baseline activity (P > 0.05, Fig E), but did reveal significant differences between clusters for dopamine activity (P < 0.0001, Fig F) and dopamine activity NBA (P < 0.0001, Fig G). We conducted Two-way ANOVA with SEX (males, females) and clusters (cluster1, cluster2, we excluded cluster3 because this group included only females) to determine if there was a SEX × cluster interaction and main effects of SEX and cluster. There were no SEX × cluster interaction (P > 0.05) for baseline activity (Fig H), dopamine activity (Fig I) and dopamine activity NBA (Fig J) and we did not need to conduct post hoc tests. Note that there were no sex differences within clusters, but there were sex differences when we compared males and females from different clusters.

The clusters were behaviorally-distinct: The F values, P values and goodness of fit (R^2^) for the lines representing the relationship between baseline activity and dopamine activity (Figure 5C) were as follows: cluster1 (F 1, 21 = 20.29, P = 0.0002, R^2^ = 0.49), cluster2 (F 1, 9 = 2.637, P = 0.1389, R^2^ = 0.23) and cluster3 (F 1, 1 = 73.61, P = 0.0739, R^2^ = 0.99), and there were no differences between clusters with respect to slopes (F 2, 31 = 1.430, P = 0.2545), but there were significant differences with respect to y-intercepts (F 2, 33 = 85.53, P < 0.0001). Similarly, the F values, P values and goodness of fit (R^2^) for the lines representing the relationship between baseline activity and dopamine activity NBA (Figure 5D) were as follows: cluster1 (F 1, 21 = 6.742, P = 0.0168, R^2^ = 0.24), cluster2 (F 1, 9 = 7.872, P = 0.0205, R^2^ = 0.47) and cluster3 (F 1, 1 = 10.05, P = 0.1945, R^2^ = 0.91), with significant differences between clusters with respect to slopes (F 2, 31 = 76.53, P < 0.0001).

The baseline activity for cluster1 (n = 23), cluster2 (n =11) and cluster3 (n = 3) were 2291 ± 301, 2371 ± 307 and 680 ± 222 cm, respectively. The dopamine activity for cluster1, 2 and 3 were 5132 ± 577, 22719 ± 2266 and 24538 ± 1583 cm, respectively. The dopamine activity NBA for cluster1, 2 and 3 were 1.34 ± 0.16, 5.27 ± 0.54 and 22.37 ± 6.92, respectively. One-way ANOVA revealed no significant differences between clusters for baseline activity (F 2, 34 = 2.237, P = 0.1223, Figure 5E), but significant differences between clusters with regards to dopamine activity (F 2, 34 = 64.04, P < 0.0001, Figure 5F) and dopamine activity NBA (F 2, 34 = 60.87, P < 0.0001, Figure 5G).

The differences between clusters could not be explained by dopamine injection site coordinates. One way ANOVA revealed the following results: AP (F 2, 34 = 0.5621, P = 0.5752), ML (F 2, 34 = 0.8245, P = 0.4470 and DV (F 2, 34 = 0.1176, P = 0.8894).

### Separation via behavioral clustering (MISSING model): Sex similarities within groups/ sex differences between groups

Having obtained the clusters, we explored SEX × cluster interaction using Two-way ANOVA with factors SEX (males, females) and clusters (cluster1-2). *We excluded cluster 3 from this specific analysis because it consisted of only females* (see Figure 5A-B). For baseline activity (Figure 5H), Two-way ANOVA did not reveal a SEX × cluster interaction (F 1, 30 = 0.3294, P = 0.5703) or a main effect of cluster (F 1, 30 = 0.5182, P = 0.4772) but did reveal a main effect of SEX (F 1, 30 = 6.676, P = 0.0149). For dopamine activity (Figure 5I), Two-way ANOVA did not reveal a SEX × cluster interaction (F 1, 30 = 2.051, P = 0.6539) or a main effect of SEX (F 1, 30 = 2.690, P = 0.1114) but did reveal a main effect of cluster (F 1, 30 = 82.50, P < 0.0001). Similarly, for dopamine activity NBA (Figure 5J), Two-way ANOVA did not reveal a SEX × cluster interaction (F 1, 30 = 2.973, P = 0.0949) or a main effect of SEX (F 1, 30 = 3.450, P = 0.0731) but did reveal a main effect of cluster (F 1, 30 = 89.72, P < 0.0001).

Comparisons between males and females in cluster1, using unpaired t-tests, revealed significant differences for baseline activity (P = 0.0208, Figure 6A), but no significant differences for dopamine activity (P = 0.0826, Figure 6B) and dopamine activity NBA (P = 0.8689, Figure 6C). Comparisons between males and females in cluster2 revealed no significant differences for baseline activity (P = 0.1504, Figure 6D), dopamine activity (P = 0.4449, Figure 6E) and dopamine activity NBA (P = 0.1752, Figure 6F). Interestingly, comparisons between males from cluster1 and females from cluster2 revealed no significant differences for baseline activity (P = 0.0846, Figure 6G), but significant differences for dopamine activity (P < 0.0001, Figure 6H) and dopamine activity NBA (P < 0.0001, Figure 6I). Likewise, comparisons between males from cluster2 and females from cluster1 revealed no significant differences for baseline activity (P = 0.1340, Figure 6J), but significant differences for dopamine activity (P = 0.0003, Figure 6K) and dopamine activity NBA (P < 0.0001, Figure 6L). Thus, sex differences appear to be driven by cluster differences not biological sex *per se*: for all variables (Figure 6), there were no sex differences *within* any cluster but there were sex differences when we compared males and females from different clusters.

**Figure 6.**
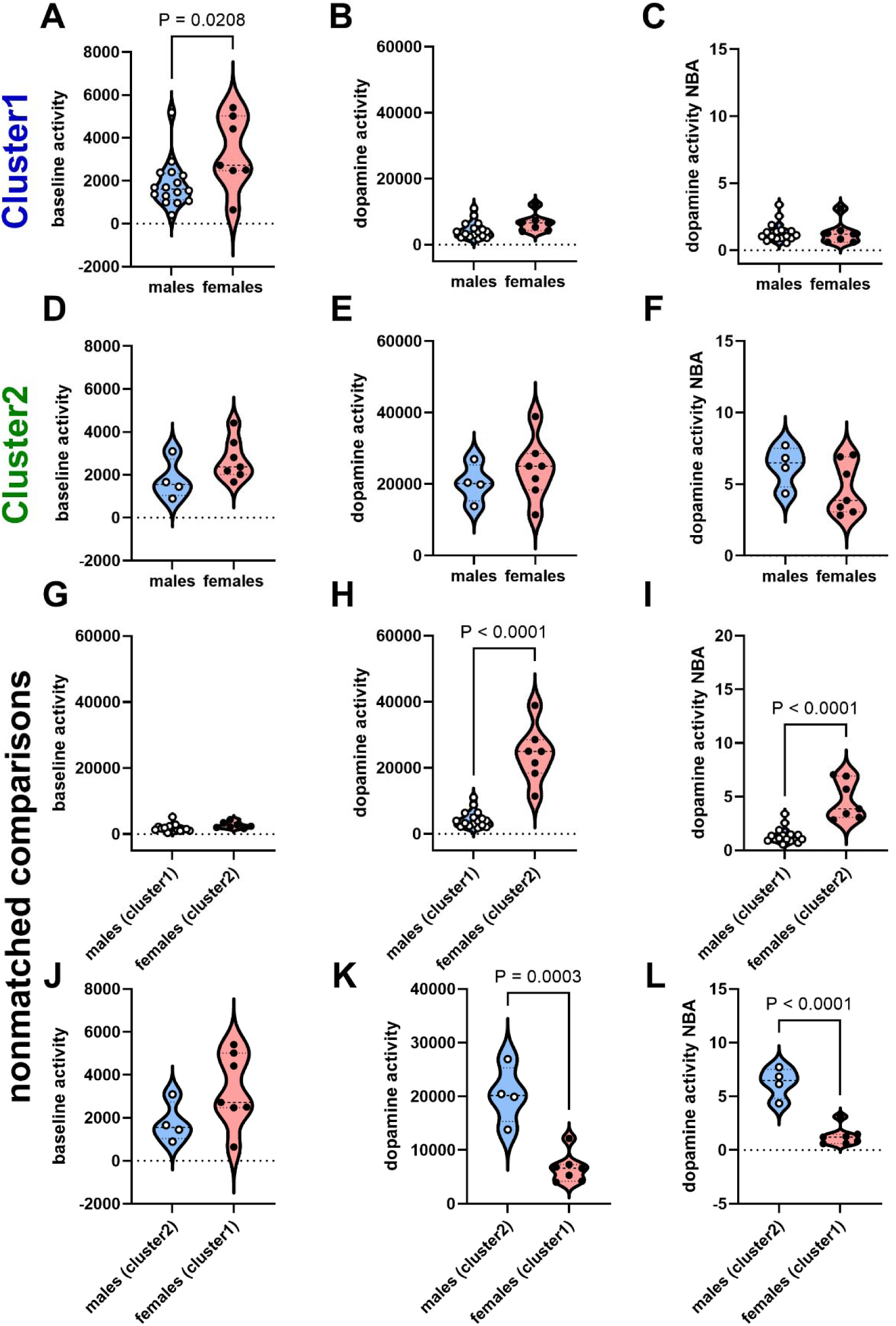
Behavioral cluster-matched and nonmatched comparisons of males and females. For cluster1 and cluster2, a Two-way ANOVA revealed no SEX × cluster interaction (see Figure 5H-J). Here, using unpaired t-tests, we compared the variables (baseline activity, dopamine activity and dopamine activity NBA) for males and females within the same group (matched comparisons) and males and females between different groups (nonmatched comparisons). Fig A-C show significant differences in baseline activity, but not in dopamine activity and dopamine activity NBA for comparisons between males and females in cluster1. Fig D-F reveal no significant differences in baseline activity, dopamine activity and dopamine activity NBA for comparisons between males and females in cluster2. Fig G-I reveal no significant differences in baseline activity, but significant differences in dopamine activity and dopamine activity NBA for comparisons between males from cluster1 and females from cluster2. Fig J-L show no significant differences in baseline activity, but significant differences in dopamine activity and dopamine activity NBA for comparisons between males from cluster2 and females from cluster1. Note the sex similarities in dopamine activity and dopamine activity NBA when we conducted matched comparisons (Fig B-C, E-F) and note that sex differences in dopamine activity and dopamine activity NBA are only observed when we conduct nonmatched comparisons (Fig H-I, K-L).

### Separation via behavioral clustering (MISSING model): Females can be more different from each other than from males

While there were no males in cluster3 (Figure 5A-B), comparisons between females in cluster3 and males in cluster1 revealed no sex differences for baseline activity (P = 0.0924, Figure 7A), but sex differences for dopamine activity (P < 0.0001, Figure 7B) and dopamine activity NBA (P < 0.0001, Figure 7C). Comparisons between females in cluster3 and males in cluster2 revealed no sex differences for baseline activity (P = 0.1194, Figure 7D) and for dopamine activity (P = 0.2713, Figure 7E), but sex differences for dopamine activity NBA (P = 0.0403, Figure 7F).

**Figure 7:**
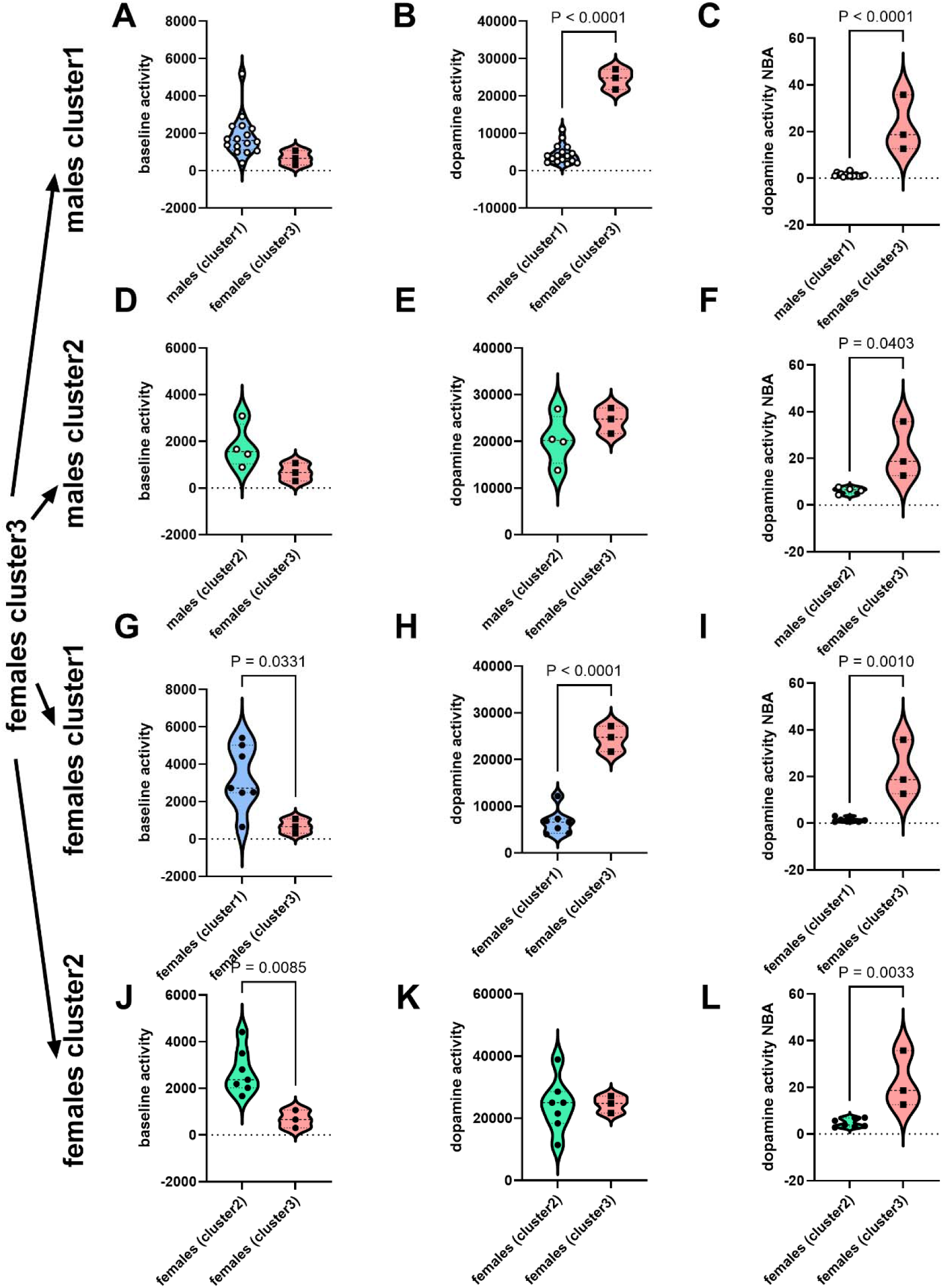
Females can be more similar to males than to other females: behavioral cluster differences exceed biological sex differences. Using unpaired t-tests, we compared the variables (baseline activity, dopamine activity and dopamine activity NBA) for males in cluster1 (Fig A-C), males in cluster2 (Fig D-F), females in cluster1 (Fig G-I) and females in cluster2 (Fig J-L) with females in cluster3 (nonmatched comparisons) Fig A-C show no significant differences in baseline activity, but significant differences in dopamine activity and dopamine activity NBA for comparisons between males in cluster1 and females in cluster3. Fig D-F reveal no significant differences in baseline activity and dopamine activity, but significant differences in dopamine activity NBA for comparisons between males in cluster2 and females in cluster3. Fig G-I reveal significant differences in baseline activity, dopamine activity and dopamine activity NBA for comparisons between females from cluster1 and females from cluster3. Fig J-L show significant differences in baseline activity and dopamine activity NBA, but no significant differences in dopamine activity for comparisons between females from cluster2 and females from cluster3. Note that females in cluster3 are more similar to males in cluster2 (Fig D-F) than to females in cluster1 (Fig G-I).

Comparisons between females in cluster3 and females in cluster1 revealed sex differences for all variables: baseline activity (P = 0.0331, Figure 7G), dopamine activity (P < 0.0001, Figure 7H) and dopamine activity NBA (P = 0.0010, Figure 7I). Comparisons between females in cluster3 and females in cluster2 revealed sex differences for baseline activity (P = 0.0085, Figure 7J) and for dopamine activity NBA (P = 0.0033, Figure 7L), but no sex differences for dopamine activity (P = 0.9371, Figure 7K). Nore that the females in cluster3 were significantly different for 3/3 variables when compared to females in cluster1 (Figure 7G, H and I), but different for only 1/3 variables when compared to males in cluster 2 (Figure 7D, E and F). *This implies that females in cluster3 were more different (behaviorally) from females in cluster1 than they were from males in cluster2*.

### Model comparisons: Dopamine NEAT analysis for groups separated by biological sex versus groups separated by behavioral clustering

We obtained the relationship between bilateral injection site coordinates (AP, ML and DV) and dopamine activity for groups 1) not separated (Figure 8A-C), 2) separated based on biological sex (Figure 8D-F), and 2) separated based on behavioral clusters (Figure 8G-I).

**Figure 8:**
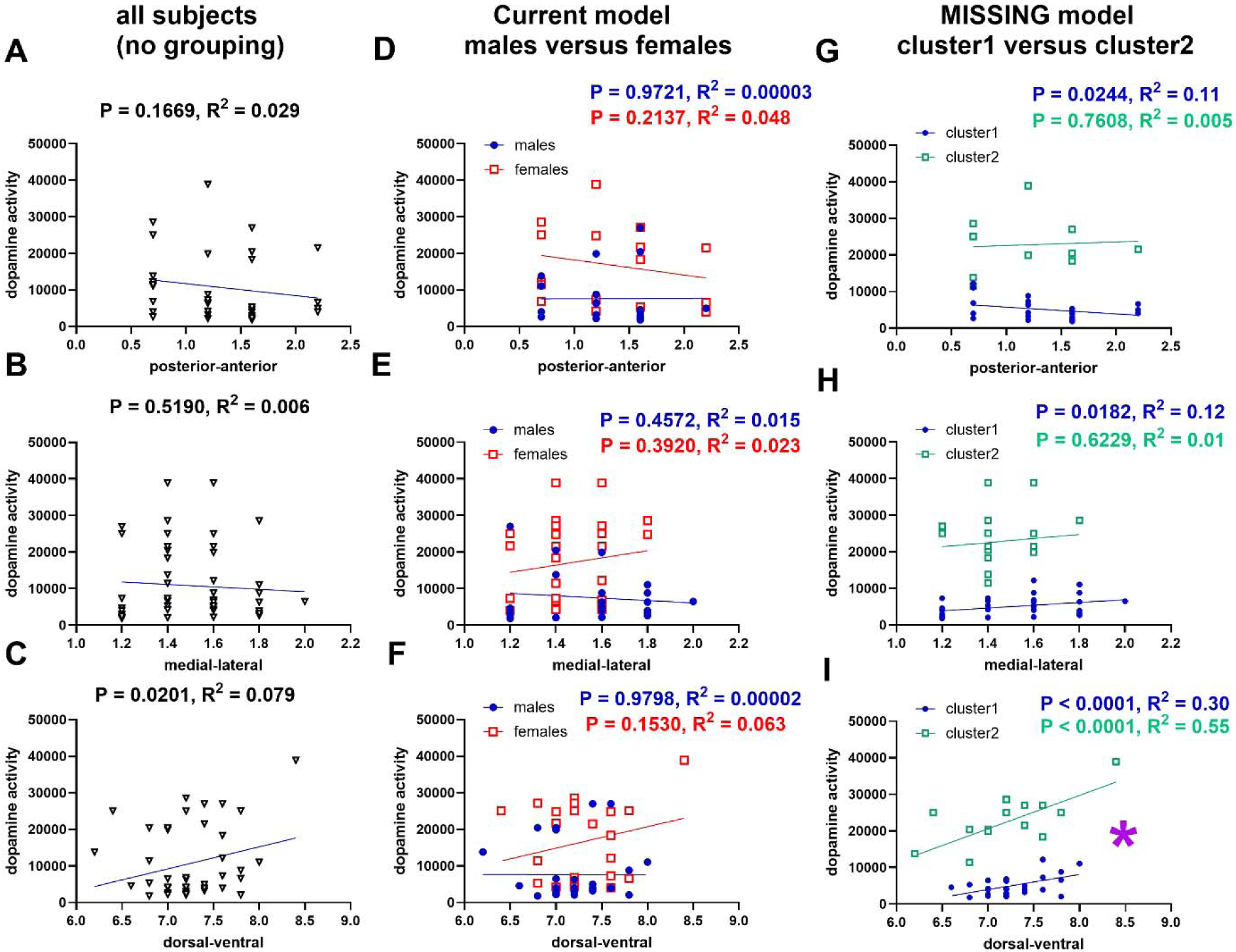
Relative to groups identified by biological sex, behavioral clusters identified by the MISSING model are more aligned with nucleus accumbens dopamine expression/activity topography: The graphs represent the relationships (from linear regression analysis) between injection site coordinates (x-axis) and dopamine activity (y-axis). The P values and R^2^ values for goodness of fit of the straight lines are shown in each graph. Fig A-C, D-F and G-I represent, respectively, all subjects (without any grouping), current model with subjects grouped by biological sex, and MISSING model with subjects grouped by behavioral cluster (for cluster1 and cluster2, we excluded cluster3 because it consisted of only 3 females and 0 males). Note that for all subjects (ungrouped), we determined that there was a significant relationship between the DV coordinates and dopamine activity (P < 0.05, Fig C). When we grouped subjects using biological sex, this relationship was no longer evident for either sex (Fig F) and there were no significant differences between males and females with regards to the relationships between DV and dopamine activity (P > 0.05, Fig F). However, when we grouped subjects using behavioral clustering (MISSING model), we detected again the relationship between DV and dopamine activity as we did for all subjects ungrouped (Fig C) but this time the relationship was more significant for both clusters (P < 0.05, Fig I) and both clusters had significantly different relationships (P < 0.05). In summary, the MISSING model revealed distinct groups (of both sexes) that were more representative of the relationship between DV and dopamine activity of the whole sample, more aligned with the expression topography of dopamine content (see first row, Table 2). By grouping according to biological sex, the groups revealed were not representative of the relationship between DV and dopamine activity of the whole sample and were not aligned with the expression topography of dopamine content in the nucleus accumbens core. Bilateral injection sites implies 2 injection sites per subject.

Ungrouped: For all subjects (ungrouped), linear regression analysis revealed no relationship between AP and dopamine activity (F 1, 66 = 1.953, P = 0.1669, R^2^ = 0.029, Figure 8A), ML and dopamine activity (F 1, 66 = 0.4203, P = 0.5190, R^2^ = 0.006, Figure 8B), but revealed a significant relationship between DV and dopamine activity (F 1, 66 = 5.674, P = 0.0201, R^2^ = 0.079, Figure 8C).

Dopamine NEAT analysis for groups separated based on biological sex – groups do not confirm NAc dopamine NEAT: For subjects grouped based on biological sex, linear regression analysis revealed (for males) no relationship between AP and dopamine activity (F 1, 38 = 0.001236, P = 0.9721, R^2^ = 0.00003, Figure 8D), ML and dopamine activity (F 1, 38 = 0.5642, P = 0.4572, R^2^ = 0.015, Figure 8E), and DV and dopamine activity (F 1, 38 = 0.0006, P = 0.9798, R^2^ = 0.000018, Figure 8F). Likewise, for females, linear regression analysis revealed no relationship between AP and dopamine activity (F 1, 32 = 1.610, P = 0.2137, R^2^ = 0.048, Figure 8D), ML and dopamine activity (F 1, 32 = 0.7529, P = 0.3920, R^2^ = 0.023, Figure 8E), and DV and dopamine activity (F 1, 32 = 2.143, P = 0.1530, R^2^ = 0.063, Figure 8F). There were no differences between males and females for the slopes of the relationship between AP and dopamine activity (F 1, 70 = 0.9420, P = 0.3351), ML and dopamine activity (F 1, 70 = 1.428, P = 0.2361) and DV and dopamine activity (F 1, 70 = 1.509, P = 0.2234).

Dopamine NEAT analysis for groups separated based on behavioral clustering – groups confirm NAc dopamine NEAT: For subjects grouped based on behavioral clustering, linear regression analysis revealed (for cluster1) relationships between AP and dopamine activity (F 1, 44 = 5.430, P = 0.0244, R^2^ = 0.11, Figure 8G), ML and dopamine activity (F 1, 44 = 6.012, P = 0.0182, R^2^ = 0.12, Figure 8H), and DV and dopamine activity (F 1, 44 = 18.73, P < 0.0001, R^2^ = 0.30, Figure 8I). For cluster2, linear regression analysis revealed no relationship between AP and dopamine activity (F 1, 20 = 0.09528, P = 0.7602, R^2^ = 0.005, Figure 8G), ML and dopamine activity (F 1, 20 = 0.2494, P = 0.6229, R^2^ = 0.01, Figure 8H), but a significant relationship between DV and dopamine activity (F 1, 20 = 24.21, P < 0.0001, R^2^ = 0.55, Figure 8I). There were no differences between cluster1 and cluster2 for the slopes of the relationship between AP and dopamine activity (F 1, 64 = 1.325, P = 0.2540) and ML and dopamine activity (F 1, 64 = 0.05465, P = 0.8159). However, there were significant differences between cluster1 and cluster2 with regards to the slope of the relationship between DV and dopamine activity (F 1, 64 = 6.887, P = 0.0108). The dopamine activity gradient (slope) for cluster1 on the AP, ML and DV (Figure 8G-I) aligns with the expression gradient of endogenous dopamine (row 1, Table 2).

## Discussion

The goal of this study was to compare the current model (which separates subjects based on biological sex) and the MISSING model (which separates subjects based on behavioral clustering) for effectiveness in identifying groups of subjects that 1) are distinct with regards to behavioral variables (baseline activity, dopamine activity and dopamine activity NBA), and 2) confirm NAc dopamine neurochemical expression/activity topography (see a summary of comparison in Table 3).

**Table 3.** Model comparison: current model versus MISSING model for sex differences.

| Current model | MISSING model |
| --- | --- |
| Separates subjects by biological sex prior to behavioral assessments (Figure 1A) | Separates subjects as individual members of behavioral groups, irrespective of biological sex, prior to assessing the impact of sex on behavioral outcomes (Figure 1B) |
| Differences in dopamine activity based on biological sex (females > males) (Figure 4C) (when all subjects were considered, there appears to be distinct groups of subjects, Figure 4D). No differences in dopamine activity when it was normalized to baseline activity (Figure 4E). | Reveals 3 behavioral clusters (Figure 5) that are distinct with regards to dopamine activity and dopamine activity normalized-to-baseline activity (NBA) and the relationship between baseline activity and dopamine activity NBA. Differences in dopamine activity only when males and females from different, not same, behavioral clusters are compared (Figure 6-7) |
| For the relationship between dorsal-ventral injection site coordinates and dopamine activity, grouping based on biological sex did not confirm NAc dopamine neurochemical expression/activity topography (Figure 8) | For the relationship between dorsal-ventral injection site coordinates and dopamine activity, grouping based on behavioral clustering confirmed NAc dopamine neurochemical expression/activity topography (Figure 8) |

The current model detected sex differences for dopamine activity (but not dopamine activity NBA), but this was complicated by our determination of more than one population of subjects in our sample. The groups identified by biological sex, males and females, were not different with regards to the relationships between injection site coordinates and dopamine activity. The MISSING model identified behavioral clusters that were significantly different from each other with regards to dopamine activity NBA. There were no sex differences in dopamine activity within the same cluster. There were sex differences in dopamine activity when males and females from different clusters were compared (nonmatched comparisons). Further, the behavioral clusters were different with regards to the relationships between dorsal-ventral injection site coordinates and dopamine activity.

Our results with intra-NAc core dopamine administration are buttressed by a previous report with intraperitoneal cocaine administration (Tigano and Job, 2024) and confirm, in line with our hypothesis, that the most distinct groups were not between sexes but between behavioral clusters. We identified a cluster that consisted of only females, which was unexpected. While we did not pick up any males for this group, this is likely due to our sample specifically. The MISSING model proposes that there will be males and females within every cluster, as revealed in a previous study (Tigano and Job, 2024).

The MISSING model proposes that 1) there exists sex similarity when we compare males and females within the same behavioral group, 2) sex differences occur when we compare males and females from different groups, and 3) even if/when we detect sex differences between males and females in the same behavioral group, these differences will not be as significant as differences between males and females from different groups. Therefore, we propose that to understand sex differences, we should compare neurochemical/biochemical differences between males and females *within* the same behavioral cluster - this would be a more rigorous way to understand sex differences that is not confounded by behavioral distinctions.

The general consensus in the field is that sex differences in psychostimulant-related behavior are driven by biological sex, hence the emphasis in NIH-funded research for SABV (Becker, 1999; Becker et al., 2016; Becker and Chartoff, 2019; Becker and Hu, 2008; Becker and Koob, 2016; Beery and Zucker, 2011; Miller et al., 2017; Prendergast et al., 2014; Shansky and Woolley, 2016; Zucker and Beery, 2019). We present evidence suggesting that SABV, as is currently applied, may not represent the most effective way to group behavioral data as there are likely to be behavioral groups more distinct from each other than males versus females.

## Acknowledgements

The author wishes to acknowledge Michael J Kuhar in whose laboratory the data was collected by MOJ. MOJ designed and conducted the behavioral experiments and statistical analysis. MOJ wrote the manuscript. The author wishes to acknowledge the support of NIH grants DA15162 and DA015040. Also this work was funded by the National Center of Research Resources P51RR165 and was supported by the Office of Research Infrastructure Programs/OD P51OD11132. This work was also supported by the Francis Lax Fund for Faculty Development at Rowan University. This work was also supported by startup funds from Rowan University, Camden, New Jersey.

## Disclosures

Dr. Martin O Job has no conflicts of interest to declare.

## Notes

### Competing Interest Statement

The authors have declared no competing interest.

